# Functional Connectivity of the Cerebellar Vermis in Bipolar Disorder and Associations with Mood

**DOI:** 10.1101/2023.02.02.526878

**Authors:** Arshaq Saleem, Gail Harmata, Shivangi Jain, Michelle W. Voss, Jess G. Fiedorowicz, Aislinn Williams, Joseph J. Shaffer, Jenny Gringer Richards, Ercole John Barsotti, Leela Sathyaputri, Samantha L. Schmitz, Gary E. Christensen, Jeffrey D. Long, Jia Xu, John A. Wemmie, Vincent A. Magnotta

## Abstract

**Purpose:** Studies of the neural underpinnings of bipolar type I disorder have focused on the emotional control network. However, there is also growing evidence for cerebellar involvement, including abnormal structure, function, and metabolism. Here, we sought to assess functional connectivity of the cerebellum with the cerebrum in bipolar disorder and to assess whether any effects might depend on mood.

**Methods:** This cross-sectional study enrolled 128 participants with bipolar type I disorder and 83 control comparison participants who completed a 3T MRI scan, which included anatomical imaging as well as resting state BOLD imaging. Functional connectivity of the cerebellar vermis to all other brain regions was assessed. Based on quality control metrics of the fMRI data, 109 participants with bipolar disorder and 79 controls were used to in the statistical analysis comparing connectivity of the vermis as well as associations with mood. Potential impacts of medications were also explored.

**Results:** Functional connectivity of the cerebellar vermis in bipolar disorder was found to differ significantly between brain regions known to be involved in the control of emotion, motor function, and language. While connections with emotion and motor control areas were significantly stronger in bipolar disorder, connection to a region associated language production was significantly weaker. In the participants with bipolar disorder, ratings of depression and mania were inversely associated with vermis functional connectivity. No effect of medications on these connections were observed.

**Conclusion:** Together the findings suggest cerebellum may play a compensatory role in bipolar disorder and when it can no longer fulfill this role, depression and mania develop. The proximity of the cerebellar vermis to the skull may make this region a potential target for treatment with transcranial magnetic stimulation.

## 1. Introduction

Bipolar type I disorder is a major psychiatric disorder with an individual lifetime prevalence of approximately 1.0%. Our prior work in bipolar disorder has observed significant differences in brain metabolism^1-4^ and function^5, 6^ in the cerebellum. Given these observations, we were interested in better understanding connectivity of the cerebellum with the cerebrum in bipolar disorder. Resting state functional connectivity employing dynamic Blood Oxygenation Level Dependent (BOLD) imaging provides the unique opportunity to study functional connectivity of neural circuits. To date there have been just a handful of studies that have employed resting state functional connectivity in bipolar disorder to study functional connectivity of the cerebellum and these studies have reported both increased and decreased functional connectivity of the cerebellum with the cerebrum^7-17^ with multiple regions involved including the vermis, dentate gyrus, crus II, and flocculus. In addition, Fetah et al. reported differential cerebellar connectivity between bipolar disorder and unipolar depression^12^. Of particular interest is the functional connectivity of the cerebellar vermis to the cerebrum due to its high degree of connectivity to nodes involved in the emotional regulation including the thalamus (vermis: VIIb and VIIIb)^18^, amygdala (vermis: IX)^19^, and anterior cingulate cortex (vermis: VI)^18^.

The hallmark feature of bipolar disorder is characterized by mood states (depression and mania) which can be severe and prolonged. A number of studies have observed metabolic^20-27^ and functional^5, 28-32^ differences associated with mood in bipolar disorder. The functional connectivity that is associated with mood remains unstudied^31, 33^. Thus, the aim of this study was to explore potential differences in bipolar type I disorder (referred to as bipolar disorder throughout) in a relatively large cohort of subjects with a focus of this work centered on the functional connectivity of the cerebellar vermis with the cerebrum.

Our initial hypothesis motivating this work was that the cerebellum serves a compensatory function to brain regions and networks involved in emotional control in bipolar disorder. Thus, we predicted that participants with bipolar disorder would have greater functional connectivity of the cerebellum vermis to regions and networks in the cerebrum involved in emotion regulation when comparing participants with bipolar disorder to controls. We further hypothesized that functional connectivity of the vermis with the cerebrum would be associated with mood such that increased functional connectivity would be related to less severe depressive and mania symptoms suggesting that the cerebellar vermis serves a compensatory role in maintaining normal mood. Finally, we explored if medications might have an association with the observed findings.

## 2. Materials and Methods

After receiving institutional review board (IRB) approval from the University of Iowa, individuals with bipolar disorder and a frequency matched comparison group were recruited into a cross-sectional neuroimaging study (**Figure 1**). Potential participants were screened and excluded from the study for comorbid neurological disorders, loss of consciousness for more than 10 minutes, current drug/alcohol abuse, or MR contraindications. Those who were eligible for the study based on their responses to the screening questions were invited to participate. Participants who provided written informed consent were enrolled into the study and evaluated using the Structured Clinical Interview for DSM Disorders to verify a psychiatric diagnosis of bipolar type I disorder. Control comparison participants were allowed to have prior diagnosis of a mood disorder such as depression as long as they were not receiving active treatment. Additional assessments included the Montgomery-Åsberg Depression Rating Scale (MADRS)^34^, Young Mania Rating Scale (YMRS)^35^, Adverse Childhood Experiences (ACEs) Questionnaire^36^, and the Columbia-Suicide Severity Rating Scale (C-SSRS)^37^. Participants also provided a list of current medications. Psychiatric medications were sorted into classes (antidepressants, antipsychotics, sedatives, anticonvulsants, stimulants, and lithium) modified from the WHO Anatomical Therapeutic Chemical System^38^. The control comparison group was identified by frequency matching for age, sex, race, and social economic status (SES) using the MacArthur Scale of Subjective Social Status^39^ of the participants with bipolar disorder. The comparison group was recruited based on advertisements in the local community and a brief screening questionnaire completed by potential participants. At the time of this analysis, the study had enrolled 128 participants with bipolar disorder and 83 control comparison participants who completed an imaging session at 3T, which included anatomical (T1, T2, and diffusion weighted), resting state functional imaging, and T1ρ imaging. In this report, we focus on the analysis of the resting state functional imaging data collected as part of this project. Most participants also completed a 7T imaging session focused on metabolic imaging as previously reported in Magnotta et al.^4^

**Figure 1.**
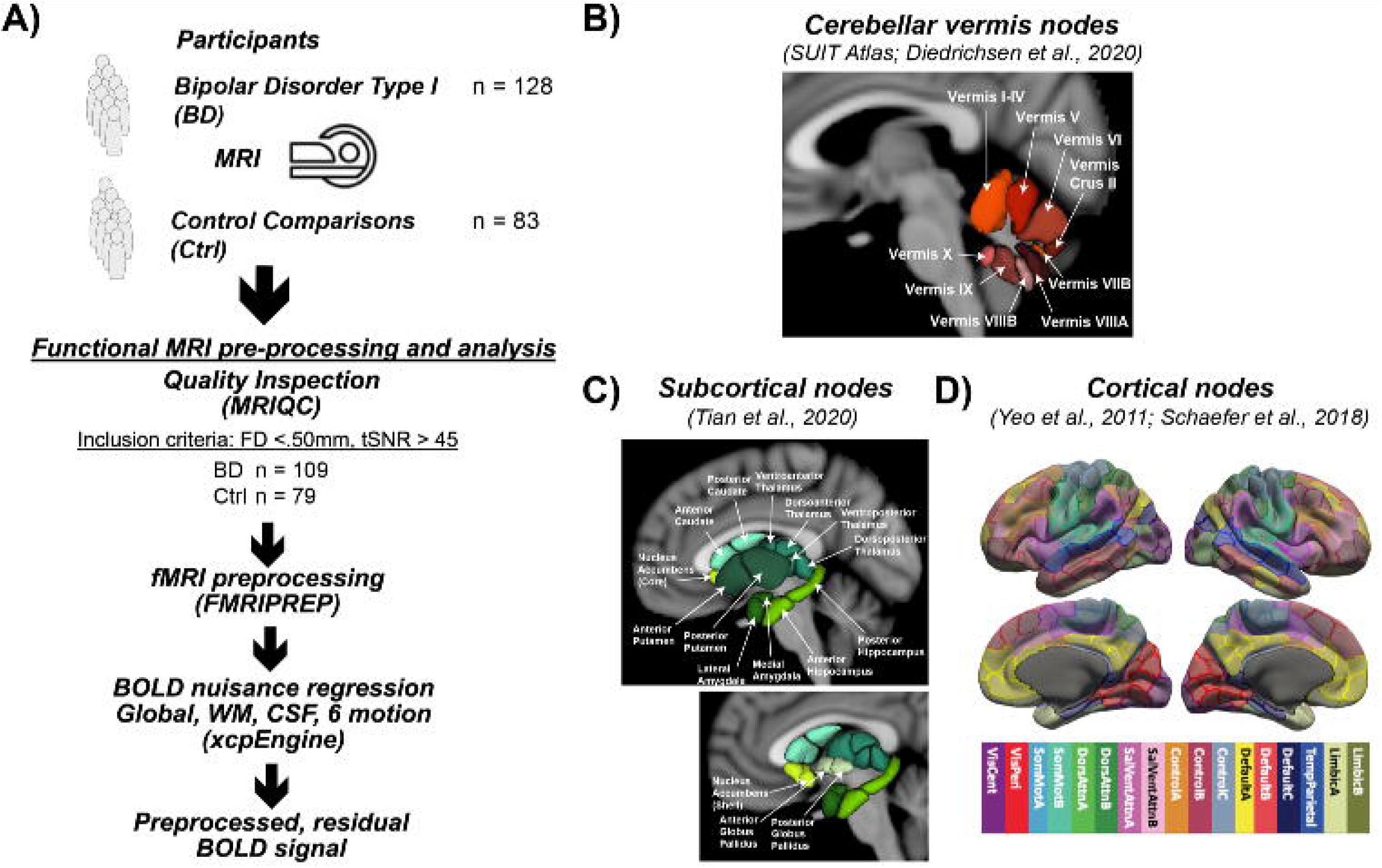
Study design. **A)** The study enrolled 128 participants with bipolar type I disorder and 83 frequency matched controls into a MR imaging study to assess differences in functional connectivity. Processing of the acquired resting state functional connectivity utilized standard image analysis pipelines including MRIQC and FMRIPREP. After preprocessing, 109 participants with bipolar disorder and 79 controls were used in the statistical analysis. **B)** Functional connectivity of the cerebellar vermis (SUIT Atlas) with the cerebellum (SUIT Atlas), subcortical (Tian Atlas), and cortical (Shaefer Atlas) regions was assessed. Specifically, the Schaefer atlas label refers to the Schaefer et al.^49^ 400 Parcels at MNI 2mm resolution, of 17 cortical networks functionally defined by Yeo et al.^84^ A detailed map of the parcellation can be seen at the atlas github page^85^. The specific image is: Schaefer2018_400Parcels_17Networks_order_FSLMNI152_2mm.nii.gz

### 2.1 MRI Data Acquisition

The MRI studies were conducted at the University of Iowa Magnetic Resonance Research Facility (MRRF) on a 3T General Electric (GE) Discovery MR750W using a 32-channel head coil which was upgraded to a 3T GE Premier MRI Scanner with a 48-channel head coil. The same imaging protocol was used across scanner configurations. A reference T1-weighted anatomical brain image was collected via a coronal fast spoiled grass sequence (TI=450ms, TE=2.0ms, TR=5.2ms, Flip Angle=12°, Matrix=256×256×220, Field of View=256×256×220mm, Bandwidth = 488 Hz/pixel) for co-registration of all functional images. Resting state functional images were acquired using an echo-planar gradient-echo sequence with a voxel size of 3.4×3.4×4mm acquired in an ascending axial slice order and no gap between slices (TE=30ms, TR=2000ms, Flip Angle = 80°, Field of View =220×220mm, Matrix=64×64×35, Bandwidth= 7812Hz/pixel; 300 volumes for a total scan time of 600sec). All participants were instructed to attend their eyes on a fixation cross, relax, and remain awake during the resting state scan.

### 2.2 MRI Data Preprocessing

All MRI data were converted from DICOM to NIFTI with the dcm2niix software. fMRIPrep (v20.2.0), a Nipype based tool, was then used to preprocess the anatomical and functional images, which included registration with the MNI atlas and nuisance regressor estimation. Several internal operations within fMRIPrep use Nilearn^40^ (version 0.6.2, RRID:SCR_001362), mostly within the functional processing workflow. For more details of the pipeline, the reader is referred to the workflows section in the fMRIPrep documentation.

#### 2.2.1 Anatomical Data Preprocessing

The T1-weighted (T1w) image was corrected for intensity non-uniformity using the N4 bias field correction algorithm^41^ (N4BiasFieldCorrection) distributed with ANTs^42^ (version 2.3.3, RRID:SCR_004757). The resulting bias field corrected T1 weighted (T1w) image was then subsequently used as the T1w-reference image throughout the remainder of the workflow. The T1w-reference image was then skull-stripped using the ANTS based brain extraction workflow (antsBrainExtraction.sh) with the OASIS30ANTs atlas used as the target template. Tissue classification was then performed on the brain-extracted T1w-reference image using FSL^43^ (fast version 5.0.9, RRID:SCR_002823) to label each voxel as cerebrospinal fluid, white-matter, or gray-matter. Next, brain surfaces were reconstructed using FreeSurfer^44^ (recon-all version 6.0.1, RRID:SCR_001847), and the brain mask was refined by employing a custom variation of the method to reconcile ANTs and FreeSurfer derived segmentations of the cortical gray-matter using Mindboggle^45^ (RRID:SCR_002438). Volume-based spatial normalization to the MNI atlas (version MNI152NLin2009cAsym) was performed through a nonlinear registration with ANTs^46^ (antsRegistration), using brain-extracted versions of both the T1w-reference and the T1 image from the MNI Atlas.

#### 2.2.2 Functional Data Preprocessing

For the 10 minutes of resting state BOLD data collected per participant, the following preprocessing steps were performed. First, a reference volume and its skull-stripped version were generated based on the median volume in the BOLD timeseries, without susceptibility distortion correction. The BOLD reference volume was then co-registered to the skull-stripped T1w reference using boundary-based registration in FreeSurfer^47^ (bbregister). Time series co-registration was then performed by concatenating transforms from 1) the rigid body realignment parameters, 2) affine and boundary-based registration of the BOLD reference image to T1w-reference image, and 3) non-linear registration of the T1w-reference image to MNI space. The resulting concatenated transform was used to transform the BOLD time series into MNI space using a single step of interpolation with a Lanczos kernel. The resulting rotation / translation parameters along with the framewise displacement (FD) were saved for use as nuisance regressors as well as quality assurance metrics respectively^48^.

Functional MRI data from participants was considered eligible for further analyses if the mean framewise displacement was below 0.50 mm, and the temporal signal-to-noise ratio (tSNR) was greater than 45. This approach resulted in 23 participants being excluded (n = 19 bipolar disorder, n = 4 controls). In all, 109 bipolar disorder participants and 79 control participants were included in the subsequent analyses evaluating the connectivity of the cerebellar vermis.

With the functional images in the MNI coordinate system, functional connectivity was assessed by combining three atlases to provide regions of interest, which included cortical brain regions from the Schaefer 400 atlas parcellated into 17 networks^49^, sub-cortical regions from the Tian atlas at Scale II^50^, and cerebellar regions from the SUIT atlas^51^. A modified version of the SUIT atlas was used in this study since only cerebellar vermal regions defining lobules VI-X are included in the atlas. To define the anterior lobe portion of the vermis (I-V), the vermal regions for this portion of the cerebellum (I-IV and V) were defined as the medial portions of these lobules with the same width as vermis VI. As a result, the full set of vermal ROIs included sub-regions I-IV, V, VI, Crus II, VIIB, VIIIA, VIIIB, IX, and X (**Figure 1B**). These regions were then subsequently used in the functional connectivity analysis as described in the following sections.

### 2.3 Functional Connectivity Analyses

#### 2.3.1 Denoising

Steps were taken to mitigate the influence of artifacts on resting state BOLD time series before assessing functional connectivity between regions. This included nuisance regression of the average signal from the participant-specific segmentations of white matter and ventricles, the global signal, as well as the 6 rigid body realignment parameters (3 translational, 3 rotational) using a modified xcpEngine design file^52^ (version 1.2.4). Temporal filtering was conducted during the nuisance regression step using AFNI^53^ (3dBandpass, Butterworth filter), which ensures BOLD fluctuations are within the frequency band of 0.01 < f < 0.08 Hz. The output of this regression based denoising is a residualized BOLD timeseries that is mean-centered at zero.

#### 2.3.2 Cerebellar Vermal Functional Connectivity

Following preprocessing, the residualized BOLD timeseries was extracted from each of the brain regions defined in the combined brain atlas described previously. The mean time series of voxels within each atlas region was used to estimate the functional connectivity by computing the Pearson’s Correlation between each vermal ROI and all ROIs included in the combined brain parcellation scheme. The resulting correlation coefficient (r) was then converted into a Fisher’s Z (ie. z(r)). Given the large number of comparisons that would result from considering the connections between all ROI pairs, we limited the connections of interest to those between the defined cerebellar vermal ROIs and all other regions. In addition, connections between the vermal ROIs that had a negative mean correlation coefficient for both groups of participants were excluded from further analyses.

#### 2.3.3 Dynamic Time Warping Analysis of Functional Connectivity

Dynamic time warping is an alternate strategy for assessing the relationship between two time series^54^. For complete details of this method, the reader is referred to Meszlényi et al.^55^ Briefly, this approach matches two time series by stretching or contracting one time series to match the second time series. In this matching, a point in time series 1 can be matched to one or more points in time series 2. In addition, the warping procedure allows multiple time points in series 1 to be matched to the same point in series 2. Dynamic time warping generates a distance metric of similarity between the time series, which is dependent on the amount of warping in both time and amplitude required to match the time series. To generate a normal distribution for further statistical analysis, dynamic time warping distance metrics for all connections in a subject are multiplied by negative one and mean centered at zero resulting in the maximum value corresponding to the greatest similarity between time series. One of the primary advantages of using this approach to analyze resting state fMRI data is that it allows for non-stationary temporal asynchrony of the two-time series. Thus, non-stationary time-lags resulting from switching states^56, 57^ as well as shape variations in the hemodynamic response between brain regions is accounted for in the analysis. For comparison purposes, the standard correlation approach calculates a correlation of two time series and strength of connection is calculated for a zero-phase lag of the two temporal signals. In addition, dynamic time warping was developed to specifically handle time series analyses, allowing it to account for autocorrelations resulting from colored noise in the data.

To implement dynamic time warping, we used R/RStudio version 4.1.1^58, 59^ with packages dtw^60^, doSNOW^61^, and tidyverse^62^. The same preprocessed data used as input into the correlation analysis described in the previous section, was also used to assess functional connectivity with dynamic time warping. We selected a maximum time lag of 8 s to allow for short temporal lags in hemodynamic response between regions.

### 2.4 Statistical Analyses

For each vermal ROI, linear regression analyses were run in R/RStudio version 4.1.1^58, 59^ with packages tidyverse^62^, broom^63^, and arsenal^64^ for all possible connections with other brain regions that survived the constraints described above. The primary analysis for this study was to compare resting state functional connectivity between participant groups (bipolar disorder versus control comparison). For this analysis, the linear models contained diagnosis as the independent variable of interest as well as covariates to control for age, sex, and tSNR with functional connectivity between the vermal ROIs and all other regions in the brain used as the dependent variables (each connection was run as a separate regression model). To account for multiple comparisons in this mass univariate approach, p-values were adjusted using false discovery rate (FDR) separately for each vermal ROI. Adjusted p-values (q-values) were considered significant at q < 0.05. However, since screening for positive connectivity resulted in a different number of statistical tests evaluated per vermal ROI, we have also reported all sites with raw uncorrected p-values < 0.001, which we refer to as trending findings. To visualize the relative contribution and direction of diagnosis to the estimated difference in functional connectivity between groups, we calculated the partial residuals using the jtools package ^65^ of the corresponding linear regression model. Finally, for all of the differences (significant and trending) connections, a follow-up analysis was conducted using the dynamic time warping data to explore the extent to which the findings were robust to analysis approach.

Secondary analyses were conducted in this study using only the participants with bipolar disorder, which focused on the association of mood or medications with the functional connectivity of the cerebellar vermis. In evaluating the association between mood and functional connectivity, MADRS or YMRS total scores were used as continuous independent variables of interest in the linear models with covariates for age, sex, and tSNR with functional connectivity between each vermal ROI and all other regions in the brain being the dependent variables. The resulting p-values were corrected for multiple comparisons using FDR. Similar linear models were used to explore associations with medications in the participants with bipolar disorder. In this analysis, linear models were used where we explored various classes of medications. Separate models were used for each class of drugs where a binary variable (0=off, 1=on) was used to code if a participant was on a particular class of medications. In these models, medication class was the variable of interest in the linear model with the same covariates used as in the other linear models. FDR correction was used to correct for multiple comparisons, which was done independently for each medication class.

For all significant/trending ROI pairs for which the target was part of the Schaefer Cortical Atlas^49^, the common name of the target was identified by matching the MNI coordinates of the ROI’s center of mass with the Harvard-Oxford Cortical Structural Atlas provided within the FSL package. As this atlas provides the percent likelihood that a given coordinate is part of a specific anatomical structure, the structure with the highest percentage match was selected as the common name reported in the figures and tables.

## 3. Results

The demographics for the participants whose resting state functional connectivity data was used in the linear models is summarized in **Table 1**. The two groups (bipolar disorder and comparison control) were well balanced for sex with each group containing approximately a 2:1 ratio of females to males. The groups had similar ages and age ranges with the mean age of the participants being 39 years of age with participants ranging from 18-70 years of age. Participants with bipolar disorder and comparison controls also had similar educational attainment of 15 years of school, but the controls did report significantly higher perceived social economic status of 1 rung on the MacArthur Scale (6.2 for comparison controls versus 5.3 for bipolar disorder). As expected, the participants with bipolar disorder had significantly higher mood ratings for both mania (YMRS total scores: 6.0 versus 0.9) and depression (MADRS total scores: 14.4 versus 2.8) as compared to the controls. They also had significantly more adverse childhood events and more suicide attempts as compared to the controls with 2 control participants having a prior suicide attempt and 50 of the participants with bipolar disorder having a prior attempt.

**Table 1.** Subject demographics

| **Variable** | **Control**<br>**(N = 79)** | **Bipolar I**<br>**(N = 109)** | **Test statistic<sup>†</sup>** | **p-value** |
| --- | --- | --- | --- | --- |
| Sex (Males, Females) | 27, 52 | 41, 68 | 0.234 | 0.628 |
| Age (years; mean +/- SD) | 39.8±14.0 | 38.5±13.2 | -0.673 | 0.502 |
| Age range (years) | 18-70 | 18-66 |  |  |
| Educational Attainment (years) | 15.4±2.09 | 15.0±2.11 | -1.347 | 0.180 |
| Self-Reported SES<sup>‡</sup> | 6.24±1.30 | 5.29±1.96 | -3.725 | <0.001* |
| ACE Score | 1.44±1.84 | 3.52±2.60 | 5.294 | <0.001* |
| YMRS Score | 0.924±1.67 | 5.97±5.99 | 7.288 | <0.001* |
| MADRS Score | 2.81±4.52 | 14.4±9.75 | 9.840 | <0.001* |
| Suicide Attempts | 0.0253±0.158 | 1.68±3.93 | 3.733 | <0.001* |

<sup>†</sup> Chi-squared test or t-test, as appropriate
<sup>‡</sup> MacArthur Scale of Subjective Social Status
\* p-value < 0.05

### 3.1 Functional Connectivity Differences Between Bipolar Disorder and Controls

When initiating this study our hypothesis was that functional connectivity between the cerebellar vermis and other brain regions, in particular regions involved in emotional regulation, would differ between participants with bipolar disorder and controls. Supporting this hypothesis, a linear regression analysis for main effect of diagnosis revealed significant differences (FDR; q < 0.05) in vermal connectivity between participants with bipolar disorder versus controls for the following three connections (see **Figure 2** and **Table 2**): vermis lobule V to left pars opercularis (Schaefer network SalVentAttnA), vermis lobule VIIIB to left postcentral gyrus (Schaefer network SomMotA), and vermis lobe VIIIB to right pre/postcentral gyrus (Schaefer network SomMotB). Additionally, trending differences (p < 0.001) in vermal connectivity were also identified between participants with bipolar disorder and controls (also shown in Figure 2 and Table 2): vermis lobule V to right posterior cingulate gyrus (Schaefer network DefaultA), vermis lobule V to left postcentral gyrus (Schaefer network SomMotA), and vermis lobule X to left lateral amygdala. With the exception of the connectivity between the vermis lobule V and left pars opercularis, the participants with bipolar disorder had a higher functional connectivity for all significant and trending differences in vermal connectivity as evident by the positive Beta coefficients estimated from the linear models.

**Table 2.** Regression analysis showing the effect of group (Bipolar-Controls) on functional connectivity (Pearson correlation) of the cerebellar vermis. All results with an uncorrected p-value < 0.001 are shown along with their false discovery rate q-value and regression beta coefficient for the effect of group. The regression analysis included covariates for age, sex, and tSNR.

| Seed Name and MNI Coordinate | Target Common Name MNI Coordinate | Atlas | Atlas Label Name (Network or Region) | p-value | q-value | Beta Coefficient |
| --- | --- | --- | --- | --- | --- | --- |
| Vermis V<br>(0, -48.5, -20) | Left Pars Opercularis<br>(-52, 8, 14) | Schaefer | SalVentAttnA | 0.0002 | 0.036* | -0.099 |
|  | Right Posterior Cingulate Gyrus<br>(6, -52, 24) | Schaefer | DefaultA | 0.0007 | 0.071 | 0.085 |
|  | Left Postcentral Gyrus<br>(-30, -38, 66) | Schaefer | SomMotA | 0.0009 | 0.071 | 0.102 |
| Vermis VIIIB<br>(0, -65, -45) | Right Pre/Postcentral Gyrus<br>(60, -6, 26) | Schaefer | SomMotB | 0.0002 | 0.040* | 0.105 |
|  | Left Postcentral Gyrus<br>(-48, -30, 58) | Schaefer | SomMotA | 0.0003 | 0.040* | 0.096 |
| Vermis X<br>(0,-48,-35) | Left Lateral Amygdala<br>(-26, -2,- 22) | Tian | Left Lateral Amygdala | 0.0005 | 0.096 | 0.079 |

\*False Discovery Rate q-value < 0.05

**Figure 2.**
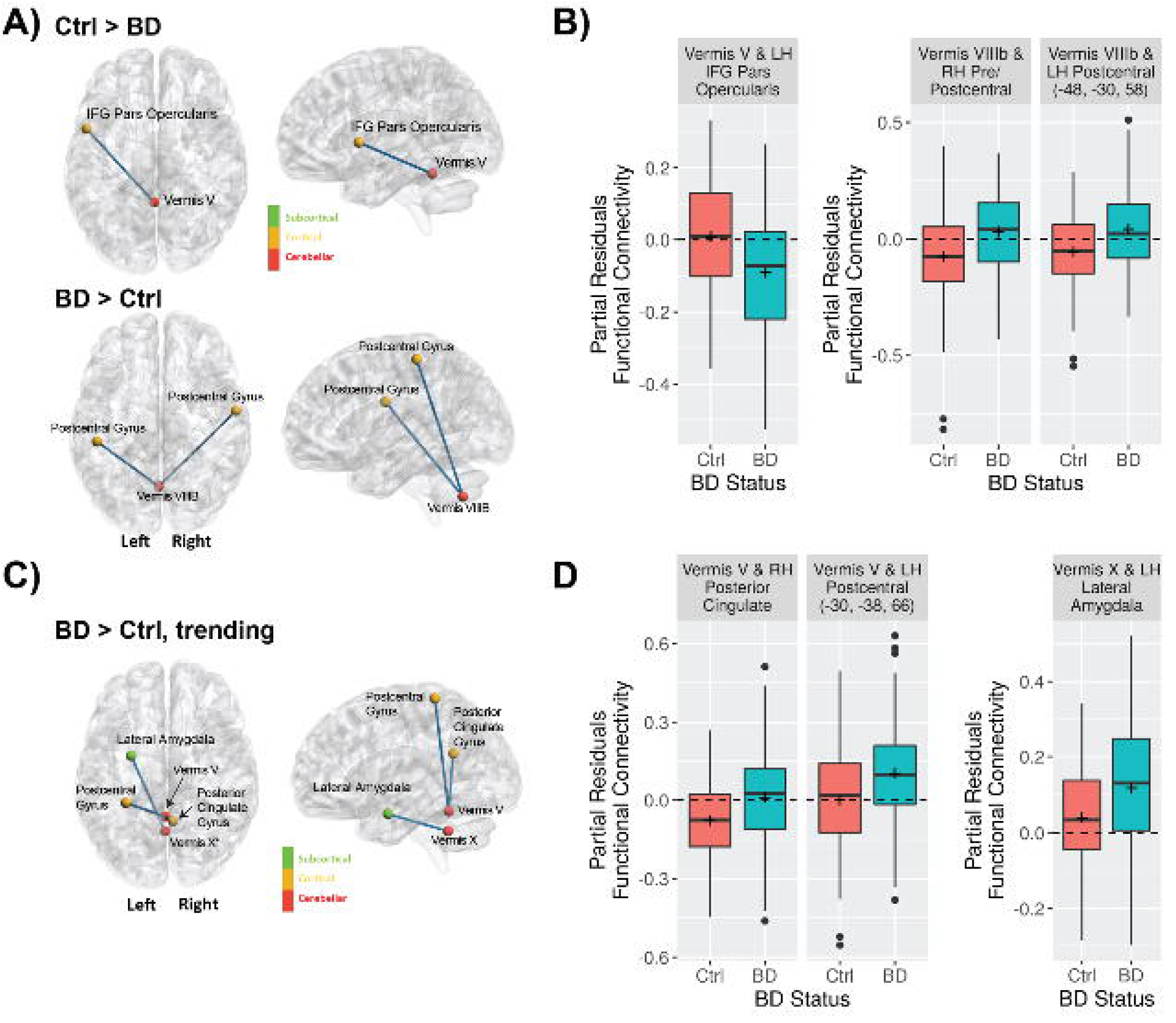
Differences in cerebellar vermal connectivity in bipolar type I disorder as compared to controls identified using a linear regression analysis including covariates for age, sex, tSNR. **A)** Shows connections with significantly (FDR corrected q-value < 0.05) different connectivity in bipolar disorder as compared to controls. **B)** Shows the functional connectivity for the bipolar and control groups for the connections shown in A. **C)** Shows connections with trending (defined as uncorrected p-value < 0.001) differences connectivity differences in bipolar disorder as compared to controls. **D)** Shows the functional connectivity for the bipolar and control groups for the connections shown in C.

#### 3.2 Dynamic Time Warping

An alternate analysis strategy, dynamic time warping, was also employed to assess the significant and trending connections of the cerebellar vermis observed using the standard time series correlation. All connections showed significant (p<0.05, uncorrected) differences between groups except the vermis lobule X to left lateral amygdala (**Supplemental Table 1**). We chose such a threshold because of the follow-up nature of the analysis. Furthermore, all connections had a beta coefficient with the same sign indicating that the differences between groups were in the same direction regardless of analysis approach. In summary all of the connections showed increased connectivity in the participants with bipolar disorder except the connection between the vermis lobule V and left pars opercularis when compared to controls.

#### 3.3 Functional Connectivity and Association with Mood in Bipolar Disorder

Evaluating the association between functional connectivity and mood may provide additional insights into how the cerebellum is involved in symptoms used to define bipolar disorder. In these analyses we evaluated the association between mood as assessed by standard scales of depression (MADRS) and mania (YMRS) with functional connectivity of the cerebellar vermis in only the participants with bipolar disorder. The significant results of the statistical models are summarized in **Figure 3** and **Table 3**. Total scores from the MADRS scale was significantly (FDR; q < 0.05) associated with functional connectivity of vermis lobule IX to left ventroposterior thalamus and vermis lobule IX to right anterior parahippocampal gyrus (Schaefer network DefaultC). In addition, there was a trending (p < 0.001) association with functional connectivity of the vermis lobule X to left ventroposterior thalamus. The direction for all of the significant and trending associations with MADRS scores was negative, such that higher functional connectivity values were associated with lower MADRS scores. Looking at the relationship between YMRS scores and functional connectivity, linear regression found a significant association (FDR; q < 0.05) between vermis lobule X and left middle frontal gyrus (Schaefer network DefaultB). As with the MADRS analysis, the direction of the association was negative, such that higher functional connectivity values were associated with lower YMRS scores.

**Table 3.**
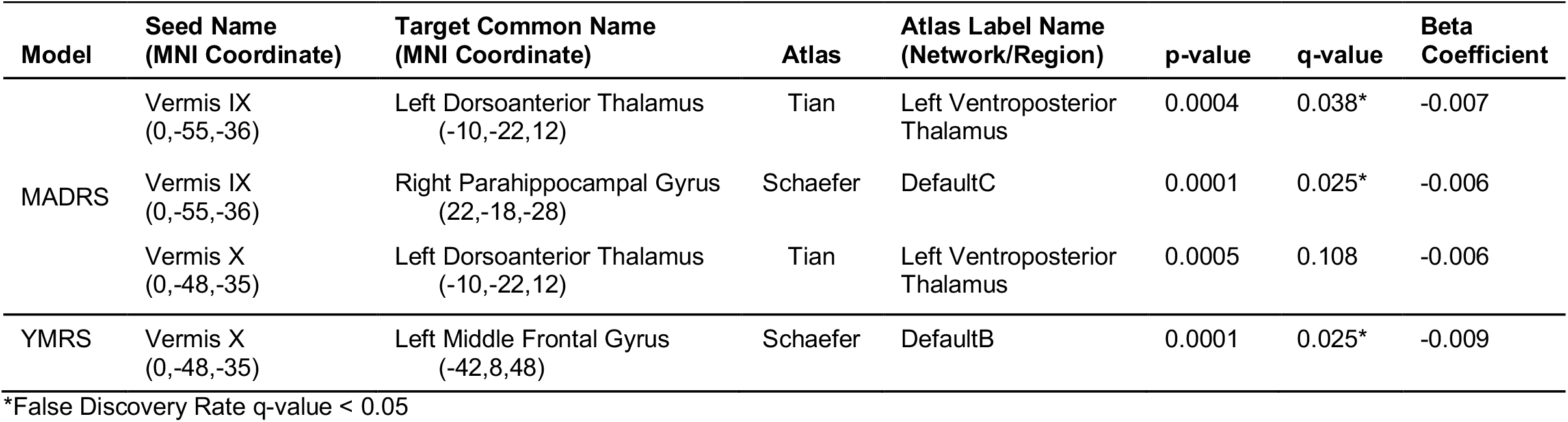
Functional connectivity of the cerebellar vermis and association with mood in bipolar disorder. All connections with a p-value < 0.001 are shown along with the corresponding FDR q-value and regression beta coefficient for the effect of mood ratings. The regression analysis included covariates for age, sex, and tSNR.

| Model | Seed Name (MNI Coordinate) | Target Common Name (MNI Coordinate) | Atlas | Atlas Label Name (Network/Region) | p-value | q-value | Beta Coefficient |
| --- | --- | --- | --- | --- | --- | --- | --- |
| MADRS | Vermis IX<br>(0,-55,-36) | Left Dorsoanterior Thalamus<br>(-10,-22,12) | Tian | Left Ventroposterior Thalamus | 0.0004 | 0.038* | -0.007 |
|  | Vermis IX<br>(0,-55,-36) | Right Parahippocampal Gyrus<br>(22,-18,-28) | Schaefer | DefaultC | 0.0001 | 0.025* | -0.006 |
|  | Vermis X<br>(0,-48,-35) | Left Dorsoanterior Thalamus<br>(-10,-22,12) | Tian | Left Ventroposterior Thalamus | 0.0005 | 0.108 | -0.006 |
| YMRS | Vermis X<br>(0,-48,-35) | Left Middle Frontal Gyrus<br>(-42,8,48) | Schaefer | DefaultB | 0.0001 | 0.025* | -0.009 |

\*False Discovery Rate q-value < 0.05

**Figure 3.**
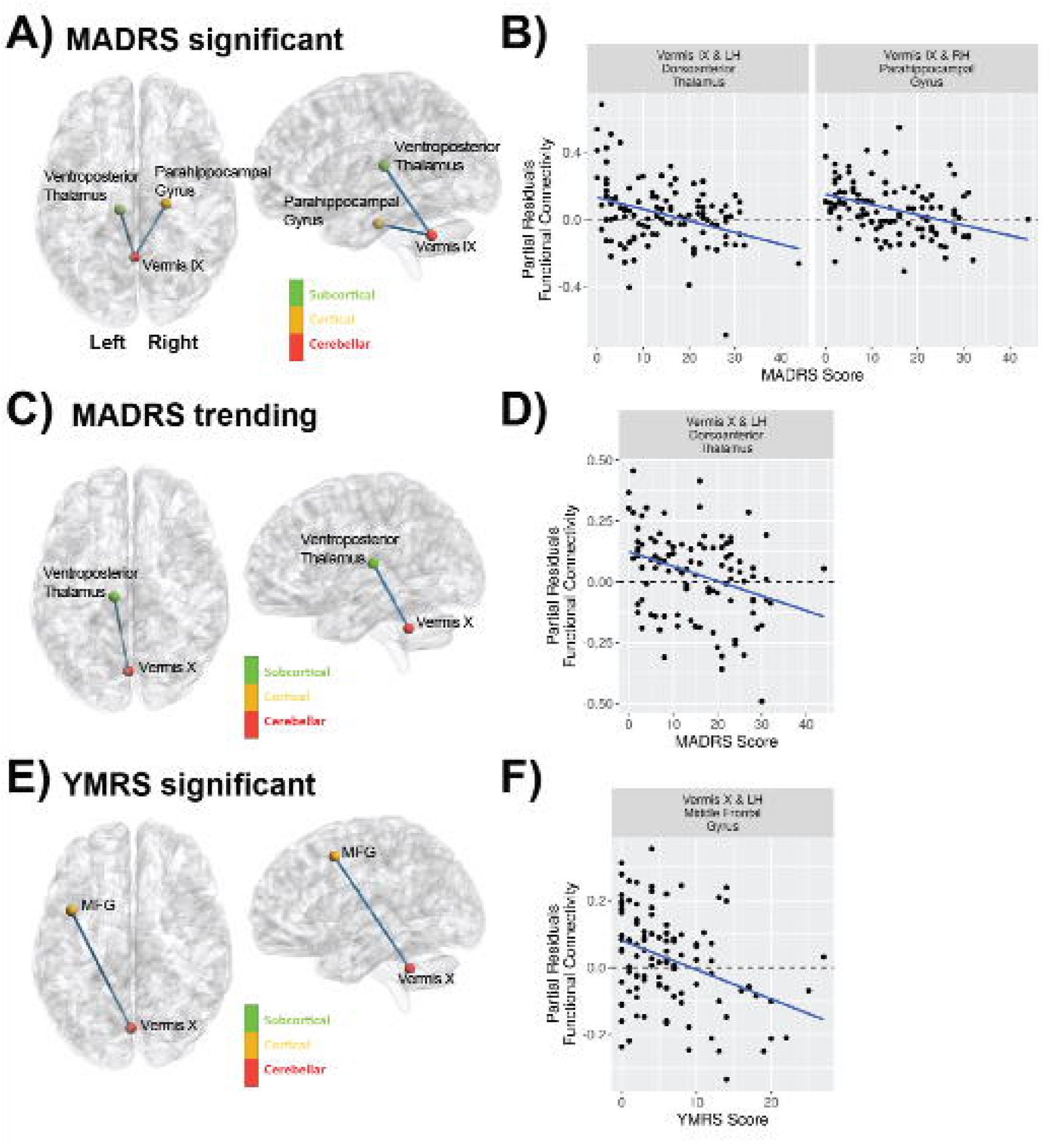
Mood as assessed with the MADRS and YMRS scales is associated with vermal connectivity in bipolar type I disorder using a regression analysis including covariates for age, sex, and tSNR. **A)** Cerebellar vermal functional connectivity with a significant (FDR corrected q-value < 0.05) association with MADRS total scores. **B)** Shows the regression between MADRS and functional connectivity for the regions identified in A. **C)** Cerebellar vermal functional connectivity with (uncorrected p-value < 0.001) association with MADRS total scores. **D)** Shows the regression between MADRS and functional connectivity for the regions identified in C. **E)** Cerebellar vermal functional connectivity with a significant (FDR corrected q-value < 0.05) association with YMRS total scores. **F)** Shows the regression between YMRS and functional connectivity for the regions identified in E.

#### 3.4 Impact of Medication on Resting State Connectivity

Linear regression models were also used to explore the effects of medication in the participants with bipolar disorder. In these analyses, medications were grouped into different classes and those with sufficient sample size defined as at least 25% of the sample size (i.e., at least 27 subjects) in each group (on and off medication) were included in separate regression models. This allowed us to explore the effects of the following medications: lithium, antipsychotics, antidepressants, anticonvulsants, and sedatives on the findings reported above. The results showed no significant effect of medication on the results even before correction for multiple comparisons (**Supplemental Table 2**). Thus, it is does not appear that medication contributed to the effects that were observed in this study. We also conducted an exploratory analysis of the effects of medication class on all of the connections of the cerebellar vermis (**Supplemental Table 3**). The resulting models found a significant effect for antidepressant medications increasing functional connectivity strength between the vermis lobule I-IV to the left posterior globus pallidus and left orbital frontal cortex (Schaefer network DefaultB). There was also a trend (p < 0.001) for lithium to increase functional connectivity between vermis lobule VIIB and the left posterior parahippocampal gyrus (Schaefer network DefaultC) as well as the vermis lobule VIIA to left precentral gyrus (Schaefer network SomMotB).

## 4. Discussion

In this study, we observed significant differences in functional connectivity of the cerebellar vermis with regions associated with emotion, language, and motor function in participants with bipolar disorder as compared to a matched control sample. Furthermore, we observed similar findings using two different approaches to assessing functional connectivity. The vermal connections where differences were observed between bipolar disorder and controls included regions involved in emotional regulation including the amygdala and posterior cingulate. Both of these connections were increased in participants with bipolar disorder as compared to the controls thus suggesting a potential compensatory role for the cerebellum associated with emotional regulation. Furthermore, vermis connectivity to regions involved in emotional salience (right anterior parahippocampal gyrus)^66, 67^ and regulation (left middle frontal gyrus)^68^ was associated with increased connectivity in participants with bipolar disorder who reported lower mood ratings. The identified regions in the cerebrum have been previously reported to be involved in processing of emotional faces^69^ as well as being abnormal when comparing the blood flow response in participants with bipolar disorder to controls using an emotional face paradigm^70-72^.

Interestingly, the vermal connections to the cerebrum associated with depression (left thalamus and right parahippocampal gyrus) were different from those associated with mania (left middle frontal gyrus). All of the relationships with mood were such that increased functional connectivity was associated with lower mood ratings. Together these findings suggests the possibility that different mood states may result from the vermis failing to serve a compensatory role with different regions of the brain associated with emotion. The reason for this is still unknown but differences in cerebellar metabolism in bipolar disorder could be one potential explanation and an observation which we have found in a sub-sample of these participants as previously reported^4^.

Connectivity of the cerebellar vermis was associated with decreased connectivity to the left pars opercularis in the participants with bipolar disorder. This is a region known to be involved in language production. Individuals with bipolar disorder especially during mania often exhibit pressured speech suggesting potential lack of inhibition of the vermis to the pars opercularis. There may also be advantages to disinhibition. The seminal work by Nancy Andreasen and colleagues on creativity and psychiatric disorders^73, 74^ suggests bipolar disorder is common in creative writers. It is tempting to speculate that a loss of inhibition from cerebellum to language areas might increase free flow of loosely related words, ideas, and concepts and thus promote creative expression.

The final set of regions with abnormal functional connectivity observed when comparing bipolar disorder to controls were sensory motor regions including precentral and postcentral gyrus. All of these findings exhibited greater connectivity between the cerebellar vermis and these sensory motor regions in bipolar disorder as compared to controls. There have been some reports of subtle abnormal motor functioning in bipolar disorder^75-79^. The data from the present study would suggest that sensory motor regions are receiving too much input from the cerebellum. In this study functional connectivity of the vermis with cortical regions was explored, but structural connectivity within these participants was not. Structural connectivity data would help to understand if these findings may result from decreased apoptosis and increased structural connectivity between these regions or if the differences were solely functional.

Development of the cerebellum starts approximately 5 weeks post conception and extends until two years postnatal also making it vulnerable to environmental perturbations both in utero as well as post-natal. One frequently reported environmental influence is adverse childhood events which occur during this critical time of brain development. As expected based on prior literature, the participants with bipolar disorder in this study exhibited a significantly greater number of adverse childhood events as compared to the controls. However, given that the measure was self-reported it is unlikely that any of the individuals were able to report events from this window of development. The importance of the vermis related to mood regulation has also been previously observed. A prior study reported that a lesion to the posterior vermis was associated with the development of mania as well as changes in functional connectivity to the prefrontal cortex, striatum, and post-central gyrus^80^. Furthermore, the vermis has been implicated in the development of psychotic features of the bipolar disorder, in particular hallucinations, using a lesion network mapping approach ^81^.

In this study we did not observe any associations between medication and the primary outcomes. However, in an exploratory analysis of this data assessing all vermal connections a significant of medication was observed for antidepressants or lithium. Each association with lithium was in the direction of increased connectivity for individuals on these classes of medications resulting in functional connectivity more similar to the comparison control sample, but this was not tested inferentially. Most of the medication studies in participants with bipolar disorder that have assessed changes in functional connectivity have been related to treatment with lithium. A prior study by Altinay et al. found that 8 weeks of treatment increased functional connectivity of the amygdala and medial orbital frontal cortex. The increased connectivity as a result of treatment provided similar connectivity as compared to a comparison control sample^82^. Another study by this same group also found that treatment with lithium normalized functional connectivity clustering coefficient by decreasing this measure during mania/hypomania and increasing this measure during depression^83^. The effects of antidepressants on resting state functional connectivity have not been studied in bipolar disorder. Nearly all of the participants with bipolar disorder in this cross-sectional study were on treatment at the time of this study with many receiving multiple classes of medication. Thus, it is difficult to fully disentangle the treatment effects especially if there are medication interactions.

While this study was one of the larger imaging studies in bipolar disorder containing 109 participants with bipolar type I disorder in the final analysis along with 79 controls, a number of limitations exist in the study. First, the study was cross-sectional in nature and the associations with mood can be only inferred based on the mood of an individual at the time they were assessed. Thus, we were not able to assess fluctuations in functional connectivity within individuals as their mood fluctuated. In the future, longitudinal studies following subjects across various mood states would help to better understand fluctuations in functional connectivity associated with mood. We also had a more limited sampling of participants with elevated mania ratings. This results from both the fact that these participants are more difficult to enroll as well as the limited ability to study these participants during the COVID-19 pandemic where significant restrictions were put on human subjects’ research and our ability of participants to leave the inpatient unit for assessment. Medication is always a challenge in studies of psychiatric disorders with most patients taking multiple classes of medications. Most of the participants in this study were on two or more medications. We did explore the effects of medication in a univariate approach and did not find any overlap with the primary findings observed in this study, however, these analyses may be more vulnerable to confounding than our primary analyses with groups balanced by age, sex and subjective SES.

In conclusion, the findings of this study support a significant role of the cerebellar vermis in bipolar disorder with abnormal functional connections between the vermis and cerebral brain regions thought to regulate mood. The strength of some of these connections further differed by mood state. Given the location and depth of the vermis relative to the skull, the vermis is potential target for treatment via transcranial magnetic stimulation. In fact, there are some on-going studies using the vermis as a target for treatment of bipolar disorder, schizophrenia, and autism. While it is unclear if the same stimulation paradigm should be used as employed for the dorsal lateral prefrontal cortex, it is potential novel therapeutic target that could help individuals with bipolar disorder to better maintain a euthymic mood state.

## Supporting information

Supplemental Materials Update

## Acknowledgements

This study was supported by funding from the National Institute of Mental Health (R01MH111578) with studies conducted on equipment (S10OD025025, S10RR028821) and facilities (UL1TR002537) supported by NIH. Some investigators received salary support from the Carver Foundation. J.A.W. was supported by NIH National Institute of Mental Health grant R01MH113325, NIH National Institute of Drug Abuse grant R01DA052953, the Roy J. Carver Charitable Trust, the Roy J. Carver Chair, a U.S. Department of Veterans Affairs Merit Review Award, and the U.S. Department of Veterans Affairs. This research was supported in part through High Performance Computing resources provided by the Information Technology Services team– Research Services at the University of Iowa, Iowa City, Iowa, directed by Joe Hetrick. We would also like to thank Joel Bruss for his help and thoughts on the use of dynamic time warping.

## Data Availability

Data from this study are available from the NIMH Data Archive (https://nda.nih.gov) under collection number 2810.

## Notes

### Competing Interest Statement

The authors have declared no competing interest.

### Summary of Updates

The supplemental material was updated to reflect the final sample used in the manuscript.

## References

1. Johnson CP, Christensen GE, Fiedorowicz JG, Mani M, Shaffer JJ, Jr., Magnotta VA, Wemmie JA. Alterations of the cerebellum and basal ganglia in bipolar disorder mood states detected by quantitative T1rho mapping. Bipolar Disord. 2018;20(4):381–90. Epub 20180107. doi: 10.1111/bdi.12581. PubMed PMID: 29316081; PMCID: PMC5995598.

2. Johnson CP, Follmer RL, Oguz I, Warren LA, Christensen GE, Fiedorowicz JG, Magnotta VA, Wemmie JA. Brain abnormalities in bipolar disorder detected by quantitative T1rho mapping. Mol Psychiatry. 2015;20(2):201–6. Epub 20150106. doi: 10.1038/mp.2014.157. PubMed PMID: 25560762; PMCID: PMC4346383.

3. Johnson CP, Follmer RL, Oguz I, Warren LA, Christensen GE, Fiedorowicz JG, Magnotta VA, Wemmie JA. Quantitative T1rho mapping links the cerebellum and lithium use in bipolar disorder. Mol Psychiatry. 2015;20(2):149. doi: 10.1038/mp.2015.10. PubMed PMID: 25727373.

4. Magnotta VA, Xu J, Fiedorowicz JG, Williams A, Shaffer J, Christensen G, Long JD, Taylor E, Sathyaputri L, Richards JG, Harmata G, Wemmie J. Metabolic abnormalities in the basal ganglia and cerebellum in bipolar disorder: A multi-modal MR study. J Affect Disord. 2022;301:390–9. Epub 20220112. doi: 10.1016/j.jad.2022.01.052. PubMed PMID: 35031333; PMCID: PMC8828710.

5. Shaffer JJ, Jr., Johnson CP, Fiedorowicz JG, Christensen GE, Wemmie JA, Magnotta VA. Impaired sensory processing measured by functional MRI in Bipolar disorder manic and depressed mood states. Brain Imaging Behav. 2018;12(3):837–47. doi: 10.1007/s11682-017-9741-8. PubMed PMID: 28674759; PMCID: PMC5752628.

6. Shaffer JJ, Jr., Johnson CP, Long JD, Fiedorowicz JG, Christensen GE, Wemmie JA, Magnotta VA. Relationship altered between functional T1rho and BOLD signals in bipolar disorder. Brain Behav. 2017;7(10):e00802. Epub 20170914. doi: 10.1002/brb3.802. PubMed PMID: 29075562; PMCID: PMC5651386.

7. Bellani M, Bontempi P, Zovetti N, Gloria Rossetti M, Perlini C, Dusi N, Squarcina L, Marinelli V, Zoccatelli G, Alessandrini F, Francesca Maria Ciceri E, Sbarbati A, Brambilla P. Resting state networks activity in euthymic bipolar disorder. Bipolar Disord. 2020;22(6):593–601. Epub 20200416. doi: 10.1111/bdi.12900. PubMed PMID: 32212391.

8. Chen G, Zhao L, Jia Y, Zhong S, Chen F, Luo X, Qiu S, Lai S, Qi Z, Huang L, Wang Y. Abnormal cerebellum-DMN regions connectivity in unmedicated bipolar II disorder. J Affect Disord. 2019;243:441–7. Epub 20180921. doi: 10.1016/j.jad.2018.09.076. PubMed PMID: 30273882.

9. Chrobak AA, Bohaterewicz B, Tereszko A, Krupa A, Sobczak A, Ceglarek A, Wielgus M, Fafrowicz M, Siwek M, Bryll A, Marek T, Dudek D. Altered functional connectivity among frontal eye fields, thalamus and cerebellum in bipolar disorder. Psychiatr Pol. 2019;54(3):487–97. Epub 20200630. doi: 10.12740/PP/OnlineFirst/104445. PubMed PMID: 33038882.

10. Cui L, Li H, Li JB, Zeng H, Zhang Y, Deng W, Zhou W, Cao L. Altered cerebellar gray matter and cerebellar-cortex resting-state functional connectivity in patients with bipolar disorder. J Affect Disord. 2022;302:50–7. Epub 20220121. doi: 10.1016/j.jad.2022.01.073. PubMed PMID: 35074460.

11. Dandash O, Yucel M, Daglas R, Pantelis C, McGorry P, Berk M, Fornito A. Differential effect of quetiapine and lithium on functional connectivity of the striatum in first episode mania. Transl Psychiatry. 2018;8(1):59. Epub 20180306. doi: 10.1038/s41398-018-0108-8. PubMed PMID: 29507281; PMCID: PMC5838223.

12. Fateh AA, Cui Q, Duan X, Yang Y, Chen Y, Li D, He Z, Chen H. Disrupted dynamic functional connectivity in right amygdalar subregions differentiates bipolar disorder from major depressive disorder. Psychiatry Res Neuroimaging. 2020;304:111149. Epub 20200717. doi: 10.1016/j.pscychresns.2020.111149. PubMed PMID: 32738725.

13. Li H, Liu H, Tang Y, Yan R, Jiang X, Fan G, Sun W. Decreased Functional Connectivity of Vermis-Ventral Prefrontal Cortex in Bipolar Disorder. Front Hum Neurosci. 2021;15:711688. Epub 20210716. doi: 10.3389/fnhum.2021.711688. PubMed PMID: 34335214; PMCID: PMC8322441.

14. Li M, Huang C, Deng W, Ma X, Han Y, Wang Q, Li Z, Guo W, Li Y, Jiang L, Lei W, Hu X, Gong Q, Merikangas KR, Palaniyappan L, Li T. Contrasting and convergent patterns of amygdala connectivity in mania and depression: a resting-state study. J Affect Disord. 2015;173:53–8. Epub 20141104. doi: 10.1016/j.jad.2014.10.044. PubMed PMID: 25462396.

15. Olivito G, Lupo M, Gragnani A, Saettoni M, Siciliano L, Pancheri C, Panfili M, Cercignani M, Bozzali M, Chiaie RD, Leggio M. Aberrant Cerebello-Cerebral Connectivity in Remitted Bipolar Patients 1 and 2: New Insight into Understanding the Cerebellar Role in Mania and Hypomania. Cerebellum. 2022;21(4):647–56. Epub 20210825. doi: 10.1007/s12311-021-01317-9. PubMed PMID: 34432230; PMCID: PMC9325834.

16. Shinn AK, Roh YS, Ravichandran CT, Baker JT, Ongur D, Cohen BM. Aberrant cerebellar connectivity in bipolar disorder with psychosis. Biol Psychiatry Cogn Neurosci Neuroimaging. 2017;2(5):438–48. doi: 10.1016/j.bpsc.2016.07.002. PubMed PMID: 28730183; PMCID: PMC5512437.

17. Wang Y, Zhong S, Jia Y, Sun Y, Wang B, Liu T, Pan J, Huang L. Disrupted Resting-State Functional Connectivity in Nonmedicated Bipolar Disorder. Radiology. 2016;280(2):529–36. Epub 20160224. doi: 10.1148/radiol.2016151641. PubMed PMID: 26909649.

18. Bernard JA, Seidler RD, Hassevoort KM, Benson BL, Welsh RC, Wiggins JL, Jaeggi SM, Buschkuehl M, Monk CS, Jonides J, Peltier SJ. Resting state cortico-cerebellar functional connectivity networks: a comparison of anatomical and self-organizing map approaches. Front Neuroanat. 2012;6:31. Epub 20120810. doi: 10.3389/fnana.2012.00031. PubMed PMID: 22907994; PMCID: PMC3415673.

19. Cauda F, Cavanna AE, D’Agata F, Sacco K, Duca S, Geminiani GC. Functional connectivity and coactivation of the nucleus accumbens: a combined functional connectivity and structure-based meta-analysis. J Cogn Neurosci. 2011;23(10):2864–77. Epub 20110125. doi: 10.1162/jocn.2011.21624. PubMed PMID: 21265603.

20. Godlewska BR, Emir UE, Masaki C, Bargiotas T, Cowen PJ. Changes in brain Glx in depressed bipolar patients treated with lamotrigine: A proton MRS study. J Affect Disord. 2019;246:418–21. Epub 20181225. doi: 10.1016/j.jad.2018.12.092. PubMed PMID: 30599363; PMCID: PMC6368663.

21. Kato T, Takahashi S, Shioiri T, Inubushi T. Brain phosphorous metabolism in depressive disorders detected by phosphorus-31 magnetic resonance spectroscopy. J Affect Disord. 1992;26(4):223–30. doi: 10.1016/0165-0327(92)90099-r. PubMed PMID: 1479134.

22. Machado-Vieira R, Zanetti MV, Otaduy MC, De Sousa RT, Soeiro-de-Souza MG, Costa AC, Carvalho AF, Leite CC, Busatto GF, Zarate CA, Jr., Gattaz WF. Increased Brain Lactate During Depressive Episodes and Reversal Effects by Lithium Monotherapy in Drug-Naive Bipolar Disorder: A 3-T 1H-MRS Study. J Clin Psychopharmacol. 2017;37(1):40–5. doi: 10.1097/JCP.0000000000000616. PubMed PMID: 27902528; PMCID: PMC5182117.

23. Moore CM, Breeze JL, Gruber SA, Babb SM, Frederick BB, Villafuerte RA, Stoll AL, Hennen J, Yurgelun-Todd DA, Cohen BM, Renshaw PF. Choline, myo-inositol and mood in bipolar disorder: a proton magnetic resonance spectroscopic imaging study of the anterior cingulate cortex. Bipolar Disord. 2000;2(3 Pt 2):207–16. doi: 10.1034/j.1399-5618.2000.20302.x. PubMed PMID: 11249799.

24. Nery FG, Weber WA, Blom TJ, Welge J, Patino LR, Strawn JR, Chu WJ, Adler CM, Komoroski RA, Strakowski SM, DelBello MP. Longitudinal proton spectroscopy study of the prefrontal cortex in youth at risk for bipolar disorder before and after their first mood episode. Bipolar Disord. 2019;21(4):330–41. Epub 20190328. doi: 10.1111/bdi.12770. PubMed PMID: 30864200.

25. Patel NC, DelBello MP, Cecil KM, Stanford KE, Adler CM, Strakowski SM. Temporal change in N-acetyl-aspartate concentrations in adolescents with bipolar depression treated with lithium. J Child Adolesc Psychopharmacol. 2008;18(2):132–9. doi: 10.1089/cap.2007.0088. PubMed PMID: 18439111.

26. Scotti-Muzzi E, Umla-Runge K, Soeiro-de-Souza MG. Anterior cingulate cortex neurometabolites in bipolar disorder are influenced by mood state and medication: A meta-analysis of (1)H-MRS studies. Eur Neuropsychopharmacol. 2021;47:62–73. Epub 20210211. doi: 10.1016/j.euroneuro.2021.01.096. PubMed PMID: 33581932.

27. Shi XF, Carlson PJ, Sung YH, Fiedler KK, Forrest LN, Hellem TL, Huber RS, Kim SE, Zuo C, Jeong EK, Renshaw PF, Kondo DG. Decreased brain PME/PDE ratio in bipolar disorder: a preliminary (31) P magnetic resonance spectroscopy study. Bipolar Disord. 2015;17(7):743–52. Epub 20151019. doi: 10.1111/bdi.12339. PubMed PMID: 26477793; PMCID: PMC5495548.

28. Brady RO, Jr., Masters GA, Mathew IT, Margolis A, Cohen BM, Ongur D, Keshavan M. State dependent cortico-amygdala circuit dysfunction in bipolar disorder. J Affect Disord. 2016;201:79–87. Epub 20160428. doi: 10.1016/j.jad.2016.04.052. PubMed PMID: 27177299; PMCID: PMC5087105.

29. Brady RO, Jr., Tandon N, Masters GA, Margolis A, Cohen BM, Keshavan M, Ongur D. Differential brain network activity across mood states in bipolar disorder. J Affect Disord. 2017;207:367–76. Epub 20161006. doi: 10.1016/j.jad.2016.09.041. PubMed PMID: 27744225; PMCID: PMC5107137.

30. Jiang W, Andreassen OA, Agartz I, Lagerberg TV, Westlye LT, Calhoun VD, Turner JA. Distinct structural brain circuits indicate mood and apathy profiles in bipolar disorder. Neuroimage Clin. 2020;26:101989. Epub 20190819. doi: 10.1016/j.nicl.2019.101989. PubMed PMID: 31451406; PMCID: PMC7229320.

31. Liu M, Wang Y, Zhang A, Yang C, Liu P, Wang J, Zhang K, Wang Y, Sun N. Altered dynamic functional connectivity across mood states in bipolar disorder. Brain Res. 2021;1750:147143. Epub 20201015. doi: 10.1016/j.brainres.2020.147143. PubMed PMID: 33068632.

32. Xiao Q, Wu Z, Hui X, Jiao Q, Zhong Y, Su L, Lu G. Manic and euthymic states in pediatric bipolar disorder patients during an emotional Go/Nogo task: A functional magnetic resonance imaging study. J Affect Disord. 2021;282:82–90. Epub 20201229. doi: 10.1016/j.jad.2020.12.105. PubMed PMID: 33401127.

33. Fan Z, Yang J, Zeng C, Xi C, Wu G, Guo S, Xue Z, Liu Z, Tao H. Bipolar Mood State Reflected in Functional Connectivity of the Hate Circuit: A Resting-State Functional Magnetic Resonance Imaging Study. Front Psychiatry. 2020;11:556126. Epub 20201027. doi: 10.3389/fpsyt.2020.556126. PubMed PMID: 33192670; PMCID: PMC7652934.

34. Montgomery SA, Asberg M. A new depression scale designed to be sensitive to change. Br J Psychiatry. 1979;134:382–9. doi: 10.1192/bjp.134.4.382. PubMed PMID: 444788.

35. Young RC, Biggs JT, Ziegler VE, Meyer DA. A rating scale for mania: reliability, validity and sensitivity. Br J Psychiatry. 1978;133:429–35. doi: 10.1192/bjp.133.5.429. PubMed PMID: 728692.

36. Felitti VJ, Anda RF, Nordenberg D, Williamson DF, Spitz AM, Edwards V, Koss MP, Marks JS. Relationship of childhood abuse and household dysfunction to many of the leading causes of death in adults. The Adverse Childhood Experiences (ACE) Study. Am J Prev Med. 1998;14(4):245–58. doi: 10.1016/s0749-3797(98)00017-8. PubMed PMID: 9635069.

37. Posner K, Brent D, Lucas C, Gould M, Stanley B, Brown G, Fisher P, Zelazny J, Burke A, Oquendo M, Mann J. Columbia-suicide severity rating scale. In: University C, editor. New York, NY2008.

38. Methodology WCCfDS. WHO Anatomical Therapeutic Chemical (ATC) Classification Oslo, Norway: WHO ATC; 2022. Available from: https://www.whocc.no/atc/structure_and_principles/.

39. Adler NE, Epel ES, Castellazzo G, Ickovics JR. Relationship of subjective and objective social status with psychological and physiological functioning: preliminary data in healthy white women. Health Psychol. 2000;19(6):586–92. doi: 10.1037//0278-6133.19.6.586. PubMed PMID: 11129362.

40. Abraham A, Pedregosa F, Eickenberg M, Gervais P, Mueller A, Kossaifi J, Gramfort A, Thirion B, Varoquaux G. Machine learning for neuroimaging with scikit-learn. Front Neuroinform. 2014;8:14. Epub 20140221. doi: 10.3389/fninf.2014.00014. PubMed PMID: 24600388; PMCID: PMC3930868.

41. Tustison NJ, Avants BB, Cook PA, Zheng Y, Egan A, Yushkevich PA, Gee JC. N4ITK: improved N3 bias correction. IEEE Trans Med Imaging. 2010;29(6):1310–20. Epub 20100408. doi: 10.1109/TMI.2010.2046908. PubMed PMID: 20378467; PMCID: PMC3071855.

42. Avants BB, Epstein CL, Grossman M, Gee JC. Symmetric diffeomorphic image registration with cross-correlation: evaluating automated labeling of elderly and neurodegenerative brain. Med Image Anal. 2008;12(1):26–41. Epub 20070623. doi: 10.1016/j.media.2007.06.004. PubMed PMID: 17659998; PMCID: PMC2276735.

43. Zhang Y, Brady M, Smith S. Segmentation of brain MR images through a hidden Markov random field model and the expectation-maximization algorithm. IEEE Trans Med Imaging. 2001;20(1):45–57. doi: 10.1109/42.906424. PubMed PMID: 11293691.

44. Dale AM, Fischl B, Sereno MI. Cortical surface-based analysis. I. Segmentation and surface reconstruction. Neuroimage. 1999;9(2):179–94. doi: 10.1006/nimg.1998.0395. PubMed PMID: 9931268.

45. Klein A, Ghosh SS, Bao FS, Giard J, Hame Y, Stavsky E, Lee N, Rossa B, Reuter M, Chaibub Neto E, Keshavan A. Mindboggling morphometry of human brains. PLoS Comput Biol. 2017;13(2):e1005350. Epub 20170223. doi: 10.1371/journal.pcbi.1005350. PubMed PMID: 28231282; PMCID: PMC5322885.

46. Fonov VS, Evans AC, McKinstry RC, Almli CR, Collins DL. Unbiased nonlinear average age-appropriate brain templates from birth to adulthood. NeuroImage. 2009;47(Supplement 1):S102. doi: 10.1016/S1053-8119(09)70884-5.

47. Greve DN, Fischl B. Accurate and robust brain image alignment using boundary-based registration. Neuroimage. 2009;48(1):63–72. Epub 20090630. doi: 10.1016/j.neuroimage.2009.06.060. PubMed PMID: 19573611; PMCID: PMC2733527.

48. Power JD, Barnes KA, Snyder AZ, Schlaggar BL, Petersen SE. Spurious but systematic correlations in functional connectivity MRI networks arise from subject motion. Neuroimage. 2012;59(3):2142–54. Epub 20111014. doi: 10.1016/j.neuroimage.2011.10.018. PubMed PMID: 22019881; PMCID: PMC3254728.

49. Schaefer A, Kong R, Gordon EM, Laumann TO, Zuo XN, Holmes AJ, Eickhoff SB, Yeo BTT. Local-Global Parcellation of the Human Cerebral Cortex from Intrinsic Functional Connectivity MRI. Cereb Cortex. 2018;28(9):3095–114. doi: 10.1093/cercor/bhx179. PubMed PMID: 28981612; PMCID: PMC6095216.

50. Tian Y, Margulies DS, Breakspear M, Zalesky A. Topographic organization of the human subcortex unveiled with functional connectivity gradients. Nat Neurosci. 2020;23(11):1421–32. Epub 20200928. doi: 10.1038/s41593-020-00711-6. PubMed PMID: 32989295.

51. Diedrichsen J, Balsters JH, Flavell J, Cussans E, Ramnani N. A probabilistic MR atlas of the human cerebellum. Neuroimage. 2009;46(1):39–46. Epub 20090205. doi: 10.1016/j.neuroimage.2009.01.045. PubMed PMID: 19457380.

52. Ciric R, Rosen AFG, Erus G, Cieslak M, Adebimpe A, Cook PA, Bassett DS, Davatzikos C, Wolf DH, Satterthwaite TD. Mitigating head motion artifact in functional connectivity MRI. Nat Protoc. 2018;13(12):2801–26. doi: 10.1038/s41596-018-0065-y. PubMed PMID: 30446748; PMCID: PMC8161527.

53. Cox RW. AFNI: software for analysis and visualization of functional magnetic resonance neuroimages. Comput Biomed Res. 1996;29(3):162–73. doi: 10.1006/cbmr.1996.0014. PubMed PMID: 8812068.

54. Sakoe H, Chiba S. Dynamic programming algorithm optimization for spoken word recognition. IEEE Transactions on Acoustics, Speech, and Signal Processing. 1978;26(1):43–9. doi: 10.1109/TASSP.1978.1163055.

55. Meszlényi RJ, Hermann P, Buza K, Gál V, Vidnyánszky Z. Resting State fMRI Functional Connectivity Analysis Using Dynamic Time Warping. Frontiers in Neuroscience. 2017;11. doi: 10.3389/fnins.2017.00075.

56. Allen EA, Damaraju E, Plis SM, Erhardt EB, Eichele T, Calhoun VD. Tracking whole-brain connectivity dynamics in the resting state. Cereb Cortex. 2014;24(3):663–76. Epub 20121111. doi: 10.1093/cercor/bhs352. PubMed PMID: 23146964; PMCID: PMC3920766.

57. Chen JE, Chang C, Greicius MD, Glover GH. Introducing co-activation pattern metrics to quantify spontaneous brain network dynamics. Neuroimage. 2015;111:476–88. Epub 20150207. doi: 10.1016/j.neuroimage.2015.01.057. PubMed PMID: 25662866; PMCID: PMC4386757.

58. Team R. RStudio: Integrated Development Environment for R. Boston, MA.: RStudio, PBC,; 2021.

59. Team RC. R: A language and environment for statistical computing. R Foundation for Statistical Computing. Vienna, Austria2021.

60. Giorgino T. Computing and Visualizing Dynamic Time Warping Alignments in R: The dtw Package. Journal of Statistical Software. 2009;31(7):1–24. doi: 10.18637/jss.v031.i07.

61. Corporation M, Weston S. doSNOW: Foreach Parallel Adaptor for the ‘snow’ Package. 2022.

62. Wickham H, Averick M, Bryan J, Chang W, McGowan L, François R, Grolemund G, Hayes A, Henry L, Hester J, Kuhn M, Pedersen T, Miller E, Bache S, Müller K, Ooms J, Robinson D, Seidel D, Spinu V, Takahashi K, Vaughan D, Wilke C, Woo K, Yutani H. Welcome to the tidyverse. Journal of Open Source Software. 2019;4(43):1686. doi: 10.21105/joss.01686.

63. Robinson D, Hayes A, Couch S. broom: Convert Statistical Objects into Tidy Tibbles. 0.8.0 ed2022.

64. Heinzen E, Sinnwell J, Atkinson E, Gunderson T, Dougherty G. An Arsenal of ‘R’ Functions for Large-Scale Statistical Summaries. 3.6.3 ed2021.

65. Long JA. jtools: Analysis and Presentation of Social Scientific Data 2022. Available from: https://cran.r-project.org/package=jtools.

66. Almeida JR, Mechelli A, Hassel S, Versace A, Kupfer DJ, Phillips ML. Abnormally increased effective connectivity between parahippocampal gyrus and ventromedial prefrontal regions during emotion labeling in bipolar disorder. Psychiatry Res. 2009;174(3):195–201. Epub 20091111. doi: 10.1016/j.pscychresns.2009.04.015. PubMed PMID: 19910166; PMCID: PMC2787954.

67. Lane RD, Reiman EM, Bradley MM, Lang PJ, Ahern GL, Davidson RJ, Schwartz GE. Neuroanatomical correlates of pleasant and unpleasant emotion. Neuropsychologia. 1997;35(11):1437–44. doi: 10.1016/s0028-3932(97)00070-5. PubMed PMID: 9352521.

68. Grecucci A, Giorgetta C, Bonini N, Sanfey AG. Reappraising social emotions: the role of inferior frontal gyrus, temporo-parietal junction and insula in interpersonal emotion regulation. Front Hum Neurosci. 2013;7:523. Epub 20130903. doi: 10.3389/fnhum.2013.00523. PubMed PMID: 24027512; PMCID: PMC3759791.

69. Fusar-Poli P, Placentino A, Carletti F, Landi P, Allen P, Surguladze S, Benedetti F, Abbamonte M, Gasparotti R, Barale F, Perez J, McGuire P, Politi P. Functional atlas of emotional faces processing: a voxel-based meta-analysis of 105 functional magnetic resonance imaging studies. J Psychiatry Neurosci. 2009;34(6):418-32. PubMed PMID: 19949718; PMCID: PMC2783433.

70. Dickstein DP, Rich BA, Roberson-Nay R, Berghorst L, Vinton D, Pine DS, Leibenluft E. Neural activation during encoding of emotional faces in pediatric bipolar disorder. Bipolar Disord. 2007;9(7):679–92. doi: 10.1111/j.1399-5618.2007.00418.x. PubMed PMID: 17988357; PMCID: PMC2946159.

71. Malhi GS, Lagopoulos J, Sachdev PS, Ivanovski B, Shnier R, Ketter T. Is a lack of disgust something to fear? A functional magnetic resonance imaging facial emotion recognition study in euthymic bipolar disorder patients. Bipolar Disord. 2007;9(4):345–57. doi: 10.1111/j.1399-5618.2007.00485.x. PubMed PMID: 17547581.

72. Sagar KA, Dahlgren MK, Gonenc A, Gruber SA. Altered affective processing in bipolar disorder: an fMRI study. J Affect Disord. 2013;150(3):1192–6. Epub 20130530. doi: 10.1016/j.jad.2013.05.019. PubMed PMID: 23726779; PMCID: PMC3922285.

73. Andreasen NC. Creativity and mental illness: prevalence rates in writers and their first-degree relatives. Am J Psychiatry. 1987;144(10):1288–92. doi: 10.1176/ajp.144.10.1288. PubMed PMID: 3499088.

74. Andreasen NC, Glick ID. Bipolar affective disorder and creativity: implications and clinical management. Compr Psychiatry. 1988;29(3):207–17. doi: 10.1016/0010-440x(88)90044-2. PubMed PMID: 3288437.

75. Bolbecker AR, Hong SL, Kent JS, Forsyth JK, Klaunig MJ, Lazar EK, O’Donnell BF, Hetrick WP. Paced finger-tapping abnormalities in bipolar disorder indicate timing dysfunction. Bipolar Disord. 2011;13(1):99–110. doi: 10.1111/j.1399-5618.2011.00895.x. PubMed PMID: 21320257; PMCID: PMC3079233.

76. Bolbecker AR, Hong SL, Kent JS, Klaunig MJ, O’Donnell BF, Hetrick WP. Postural control in bipolar disorder: increased sway area and decreased dynamical complexity. PLoS One. 2011;6(5):e19824. Epub 20110518. doi: 10.1371/journal.pone.0019824. PubMed PMID: 21611126; PMCID: PMC3097205.

77. Bolbecker AR, Mehta C, Johannesen JK, Edwards CR, O’Donnell BF, Shekhar A, Nurnberger JI, Steinmetz JE, Hetrick WP. Eyeblink conditioning anomalies in bipolar disorder suggest cerebellar dysfunction. Bipolar Disord. 2009;11(1):19–32. doi: 10.1111/j.1399-5618.2008.00642.x. PubMed PMID: 19133963.

78. Chrobak AA, Siuda-Krzywicka K, Siwek GP, Arciszewska A, Siwek M, Starowicz-Filip A, Dudek D. Implicit motor learning in bipolar disorder. J Affect Disord. 2015;174:250–6. Epub 20141209. doi: 10.1016/j.jad.2014.11.043. PubMed PMID: 25527995.

79. Sasayama D, Hori H, Teraishi T, Hattori K, Ota M, Matsuo J, Kawamoto Y, Kinoshita Y, Hashikura M, Amano N, Higuchi T, Kunugi H. More severe impairment of manual dexterity in bipolar disorder compared to unipolar major depression. J Affect Disord. 2012;136(3):1047–52. Epub 20111212. doi: 10.1016/j.jad.2011.11.031. PubMed PMID: 22169250.

80. Lupo M, Olivito G, Siciliano L, Masciullo M, Molinari M, Cercignani M, Bozzali M, Leggio M. Evidence of Cerebellar Involvement in the Onset of a Manic State. Front Neurol. 2018;9:774. Epub 20180912. doi: 10.3389/fneur.2018.00774. PubMed PMID: 30258401; PMCID: PMC6143664.

81. Kim NY, Hsu J, Talmasov D, Joutsa J, Soussand L, Wu O, Rost NS, Morenas-Rodriguez E, Marti-Fabregas J, Pascual-Leone A, Corlett PR, Fox MD. Lesions causing hallucinations localize to one common brain network. Mol Psychiatry. 2021;26(4):1299–309. Epub 20191028. doi: 10.1038/s41380-019-0565-3. PubMed PMID: 31659272.

82. Altinay M, Karne H, Anand A. Lithium monotherapy associated clinical improvement effects on amygdala-ventromedial prefrontal cortex resting state connectivity in bipolar disorder. J Affect Disord. 2018;225:4–12. Epub 20170627. doi: 10.1016/j.jad.2017.06.047. PubMed PMID: 28772145; PMCID: PMC5844774.

83. Spielberg JM, Matyi MA, Karne H, Anand A. Lithium monotherapy associated longitudinal effects on resting state brain networks in clinical treatment of bipolar disorder. Bipolar Disord. 2019;21(4):361–71. Epub 20181214. doi: 10.1111/bdi.12718. PubMed PMID: 30421491; PMCID: PMC8593846.

84. Yeo BT, Krienen FM, Sepulcre J, Sabuncu MR, Lashkari D, Hollinshead M, Roffman JL, Smoller JW, Zollei L, Polimeni JR, Fischl B, Liu H, Buckner RL. The organization of the human cerebral cortex estimated by intrinsic functional connectivity. J Neurophysiol. 2011;106(3):1125–65. Epub 20110608. doi: 10.1152/jn.00338.2011. PubMed PMID: 21653723; PMCID: PMC3174820.

85. Yeo T. Available from: https://github.com/ThomasYeoLab/CBIG/tree/master/stable_projects/brain_parcellation/Schaefer2018_LocalGlobal/Parcellations/MNI.

