## Supplemental Materials Update for "Functional Connectivity of the Cerebellar Vermis in Bipolar Disorder and Associations with Mood"

**Supplemental Table 1. Regression analysis of the dynamic time warping distance metric for all connections with a p-value < 0.001 for the effect of group (Bipolar-Control) from the primary analysis using Pearson correlations. Covariates for age, sex, and tSNR were included in the model.**

| Seed Name and MNI Coordinate | Target Common Name and MNI Coordinate | Atlas | Atlas Label Name (Network or Region) | p-value | Beta Coefficient |
|---|---|---|---|---|---|
| Vermis V (0, -48.5, -20) | Left IFG Pars Opercularis (-52, 8, 14) | Schaefer | SalVentAttnA | 0.0018 | -0.353 |
| | Right Posterior Cingulate Gyrus (6, -52, 24) | Schaefer | DefaultA | 0.0001 | 0.489 |
| | Left Postcentral Gyrus (-30, -38, 66) | Schaefer | SomMotA | 0.0227 | 0.270 |
| Vermis VIIIB (0, -65, -45) | Right Pre/Postcentral Gyrus (60, -6, 26) | Schaefer | SomMotB | 0.0098 | 0.352 |
| | Left Postcentral Gyrus (-48, -30, 58) | Schaefer | SomMotA | 0.0038 | 0.414 |
| Vermis X (0,-48,-35) | Left Lateral Amygdala (-26, -2,- 22) | Tian | Left Lateral Amygdala | 0.0643 | 0.210 |

**Supplemental Table 2. Regression analysis evaluating functional connectivity in participants with bipolar disorder on and off various classes of medications. Only regions identified with a p-value < 0.001 for the effect of group (Bipolar-Control) from the primary analysis using Pearson correlations were examined. Each connection and medication class were assessed separately. Covariates for age, sex, and tSNR were included. Medication class did not contribute significantly to the observed functional connectivity for these connections in the participants with bipolar disorder.**

| Seed Name (MNI Coordinate) | Target Common Name (MNI Coordinate) | Medication Class | p-value | Beta Value |
|---|---|---|---|---|
| Vermis Lobule V (0, -48.5, -20) | Left Pars Opercularis (-52, 8, 14) | Antidepressants | 0.225 | -0.0458 |
| | | Antipsychotics | 0.956 | 0.00183 |
| | | Sedatives | 0.353 | -0.0333 |
| | | Anticonvulsants | 0.178 | -0.0475 |
| | | Lithium | 0.316 | -0.0369 |
| | Right Posterior Cingulate Gyrus (6, -52, 24) | Antidepressants | 0.148 | 0.0562 |
| | | Antipsychotics | 0.132 | -0.0516 |
| | | Sedatives | 0.812 | 0.00882 |
| | | Anticonvulsants | 0.829 | -0.00791 |
| | | Lithium | 0.583 | 0.0209 |
| | Left Postcentral Gyrus (-30, -38, 66) | Antidepressants | 0.354 | -0.0389 |
| | | Antipsychotics | 0.930 | 0.00324 |
| | | Sedatives | 0.446 | 0.0305 |
| | | Anticonvulsants | 0.110 | 0.0625 |
| | | Lithium | 0.516 | -0.0267 |
| Vermis Lobule VIIIB (0, -65, -45) | Right Pre/Postcentral Gyrus (60, -6, 26) | Antidepressants | 0.150 | 0.0535 |
| | | Antipsychotics | 0.271 | -0.0361 |
| | | Sedatives | 0.974 | 0.00117 |
| | | Anticonvulsants | 0.561 | -0.0203 |
| | | Lithium | 0.758 | 0.0112 |
| | Left Postcentral Gyrus (-48, -30, 58) | Antidepressants | 0.902 | -0.00479 |
| | | Antipsychotics | 0.765 | 0.0102 |
| | | Sedatives | 0.936 | 0.00297 |

| | | | | |
|---|---|---|---|---|
| | | Anticonvulsants | 0.648 | 0.0166 |
| | | Lithium | 0.552 | 0.0225 |
| Vermis Lobule X (0,-48,-35) | Left Lateral Amygdala (-26, -2,- 22) | Antidepressants | 0.141 | -0.0527 |
| | | Antipsychotics | 0.301 | 0.0327 |
| | | Sedatives | 0.497 | 0.0232 |
| | | Anticonvulsants | 0.611 | 0.0171 |
| | | Lithium | 0.591 | -0.0188 |

**Supplemental Table 3. Regression analysis evaluating functional connectivity in participants with bipolar disorder on and off various classes of medications evaluating the effect of all cerebellar vermis connections. Covariates for age, sex, and tSNR were included in the model. All cerebellar vermis connections with a p-value < 0.001 for the effect of medication class are shown along with the corresponding FDR corrected q-value and estimated beta coefficient for the effect of medication.**

| Medication Class | Seed Name and MNI Coordinate | Target Common Name and MNI Coordinate | Atlas | Atlas Label Name (Network or Region) | p-value | q-value | Beta Value |
|---|---|---|---|---|---|---|---|
| Antidepressants | Vermis I-IV (0,-45.5,-16.5) | Left Posterior Globus (-22,-8,-2) | Tian | Left Posterior Globus Pallidus | 0.0003 | 0.048* | 0.145 |
| | Vermis I-IV (0,-45.5,-16.5) | Left Orbitofrontal Cortex (-36,22,-16) | Schaefer | DefaultB | 0.0005 | 0.048* | 0.139 |
| Antipsychotics | -- | -- | -- | -- | -- | -- | -- |
| Sedatives | -- | -- | -- | -- | -- | -- | -- |
| Anticonvulsants | -- | -- | -- | -- | -- | -- | -- |
| Lithium | Vermis VIIb (1,-69,-31) | Left Posterior Parahippocampal Gyrus (-30,-32,-18) | Schaefer | DefaultC | 0.0005 | 0.103 | 0.122 |
| | Vermis VIIIa (1,-70,-42) | Left Precentral Gyrus (-60,-2,10) | Schaefer | SomMotB | 0.0007 | 0.190 | 0.136 |
